# Spatiotemporal Abstraction Theory: Re‐Interpretation of Localized Cortical Networks

**DOI:** 10.1101/2025.07.07.663596

**Authors:** Gene Fridman

**Affiliations:** Department of Otolaryngology, Johns Hopkins University, Baltimore, MD, USA; Department of Biomedical Engineering, Johns Hopkins University, Baltimore, MD, USA

## Abstract

The brain excels at extracting meaning from noisy and degraded input, yet the computational principles that underlie this robustness remain unclear. We propose a theory of spatiotemporal abstraction (STA), in which localized cortical networks integrate inputs across space and time to produce multi-scale, concept-level representations that remain stable despite loss of detail. We demonstrate how this principle explains a long-standing paradox of how cochlear implant patients can understand speech despite severely scrambled neural patterns. STA provides a unified framework that explains fundamental questions: Why do we have so many neurons that respond very similarly in one cortical location? Why do we have different inhibitory neurons? It also forces us to re-examine long-standing explanations of memory, creativity, illusions, attractor dynamics, excitatory-to-inhibitory balance, and the structure and purpose of the ubiquitous canonical circuits seen throughout the brain. We conclude with STA implications for improving neural implants and artificial neural networks.

## Introduction

Cochlear implants (CIs) have enabled profoundly deaf individuals to recognize speech since their introduction in the early 1970s. At the time, the concept was controversial: the electrical stimulation delivered by these devices would grossly distort the carefully orchestrated neural activity found in normal hearing. How such distorted input could be interpreted by cortical circuits adapted to clean, structured sensory input remained an open question. Yet despite this mismatch, CI users can understand sentences with remarkable success even though the neural signals reaching their auditory cortex are sparse, noisy, and highly degraded ^1,2^.

This paradox highlights a broader principle: the brain is remarkably adept at extracting meaning from incomplete, ambiguous, or distorted input. This capacity for abstraction underlies not only speech perception, but also object recognition, memory, language, and decision making^3,4^. Hierarchical organization and functional invariance have been extensively documented across sensory and association cortices. The explanation of how the brain can maintain memory, understanding, and generalize in presence of noise has been challenging and primarily thought to be hidden in the depths of synaptic weights accessible primarily through thorough careful computational modeling.^5^

Here we propose a theory of **spatiotemporal abstraction (STA)**, in which conceptual meaning arises through systematic integration of neural activity across space and time within each localized cortical module. In this framework, concepts are encoded as broad, distributed patterns of activity, while fine-scale details encode specific instances. The degree of abstraction depends on the scale of integration: as spatial and temporal kernels widen, representations lose detail but gain invariance.

To illustrate this principle, we examine spectrograms of spoken sentences subjected to progressive spatial blurring and compare them to quantized spectrograms that simulate CI processing. As blur increases, the spectrograms increasingly resemble their quantized counterparts—mirroring how implant users comprehend speech despite massive information loss. This convergence suggests that abstraction is not a high-level symbolic operation, but a computational consequence of hierarchical spatiotemporal integration in neural systems.

This observation motivates a general theory of how the brain encodes meaning across sensory and cognitive domains. We formalize this idea through a model of spatiotemporal abstraction, in which neural representations are shaped by the scale of integration across space and time. As integration widens, input patterns are blurred, specific details are lost, and more generalized, concept-level structure emerges. We develop this framework mathematically and demonstrate its implications using speech spectrograms, showing how coarse integration leads to convergence with cochlear implant representations. We further propose that this principle of emergent abstraction through hierarchical integration applies broadly across sensory modalities and brain regions, offering a unified account of how meaning survives noise.

### Spatiotemporal Abstraction (STA) Model

We model a cortical processing module as a population of spatiotemporal integrators that vary in their integration scale across both space and time. While this model generalizes across neural systems, it is most intuitively illustrated by the retina, where many of its core features are anatomically and functionally explicit.

The retina is known to have extensive computational capability, that complements that of cortical processing.^6^ Retinal ganglion cells (RGCs) form mosaics of receptive fields that vary in size and temporal dynamics, effectively tiling the visual field with overlapping layers of integration. Each RGC computes a spatiotemporal average of the photoreceptor input within its receptive field. Cells with small receptive fields capture fine detail; those with larger receptive fields produce blurred, more abstract representations. Crucially, multiple mosaics of differing spatial scales operate in parallel, allowing the same visual scene to be simultaneously encoded at different levels of abstraction ^7–10^.

We propose that similar multi-scale integration architectures exist throughout the cortex, where they have often gone unrecognized as mechanisms of abstraction. To formalize this idea, we define the input to a generic neural network as a function *I(s,t)*, where *s* represents spatial position or, more generally, a distribution over strongly weighted synaptic inputs, and *t* is time. For each output unit in the network, we define a spatiotemporal integration kernel:

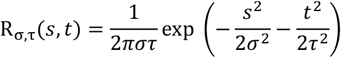

Here, *σ* and *τ* determine the spatial and temporal scales of integration, respectively. The output of a cell at a given integration scale is computed via convolution:

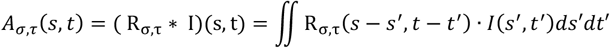

This operation filters the input through a spatiotemporal lens whose scale determines the level of abstraction. At small *σ* and *τ*, the output closely reflects local input detail; at larger scales, fine-grained features are smoothed away, leaving a more generalized encoding (**Fig. 1**). Conceptually, this defines a continuum from instance-level specificity to category-level abstraction.

**Figure 1.**
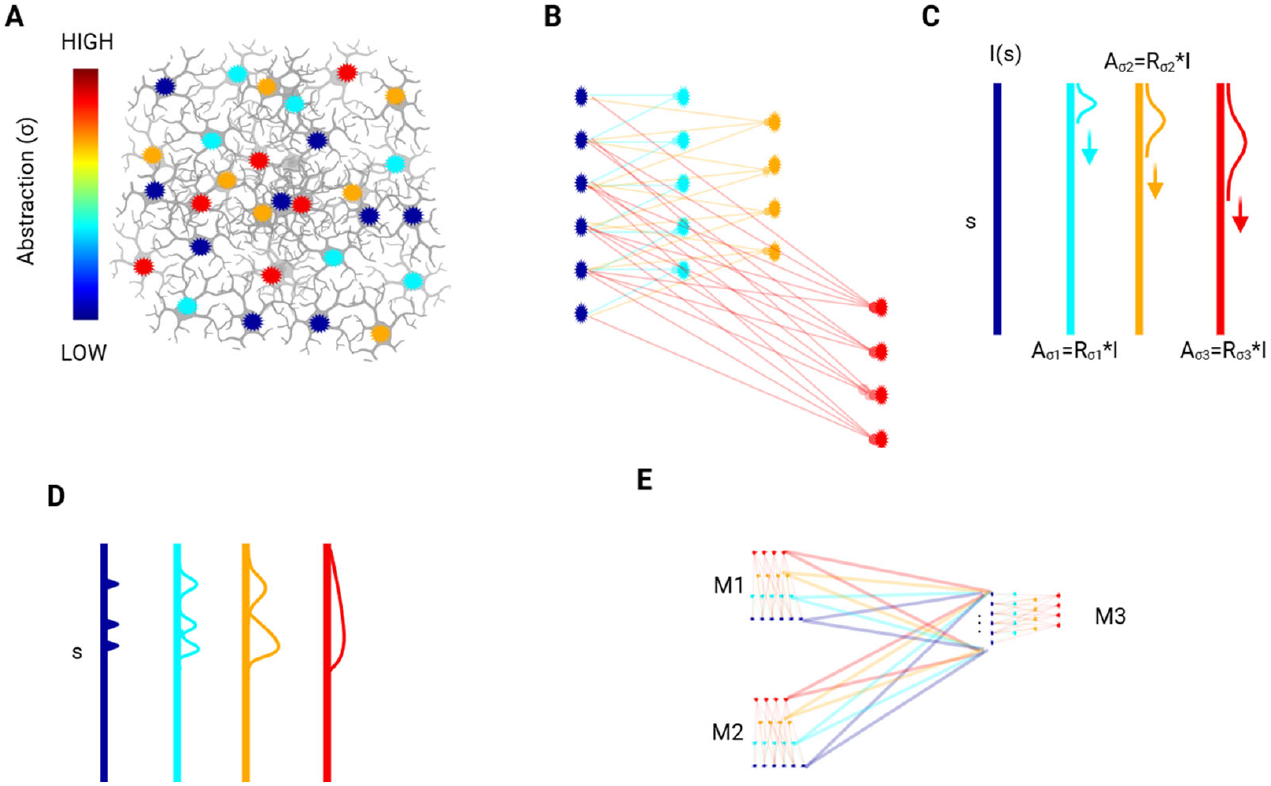
Illustration of the spatiotemporal abstraction (STA) model, shown here in its spatial form for simplicity. (**A**) Schematic of a neural module composed of interconnected units (e.g., neurons or subnetworks), color-coded by their spatial integration scale (σ). Units with larger σ integrate over broader subsets of the input population and thus encode more abstract features. (**B**) Abstracted hierarchy corresponding to panel A, where each layer represents a different spatial scale of integration applied directly to the input layer. Higher-σ units pool from more distant inputs but do not operate on each other. Red neurons are offset to illustrate the integration windows. (**C**) Mathematical formulation of STA: input signal I(s) is convolved with a spatial Gaussian kernel R_σ_(s), producing output A_σ_(s) at each abstraction level. (**D**) Example of abstraction via Gaussian convolution: a structured input (blue) is progressively smoothed into coarse representations at higher σ, illustrating the tradeoff between resolution and abstraction. (**E**) Conceptual diagram of cross-module abstraction: two modules (M1, M2), each containing their own abstraction hierarchies, project to a downstream module (M3). M3 integrates across abstraction levels and domains, supporting generalization and concept formation.

In **Figure 1A–C**, we illustrate this principle schematically. **Figure 1A** shows a network receiving input from the input level units in blue, with the higher-level integrators color-coded by weight according to their level of abstraction. **Figure 1B** redistributes the units to show synaptic relationships, emphasizes the strongest connections that define a cell’s functional receptive field, while **Figure 1C** abstracts this pattern as a Gaussian kernel. As shown in **Figure 1D**, increasing the spatial and temporal integration scale yields a progressive transformation from detailed representations to increasingly abstract ones. **Figure 1E** indicates that the connections between modules allows for information from each abstraction layer to be incorporated into other modules’ inputs.

### Application to Cochlear Implant Perception

To illustrate how neural circuits can benefit from spatiotemporal abstraction, we examine a long-standing puzzle in auditory neuroscience: how cochlear implant (CI) users can understand speech despite severely distorted neural input ^1,2,11^. CIs restore hearing by electrically stimulating auditory nerve fibers at different points along the cochlea’s tonotopic axis. The stimulation is typically delivered as a sequence of pulses across electrode contacts, each representing a coarse frequency band ^1,12^.

This stimulation strategy introduces two major forms of distortion that we simulate here ^12,13^. First, pulsatile stimulation disrupts the natural fine structure of the acoustic signal, generating phase-locked neural responses that replace the detailed temporal patterns found in normal hearing. In this case we simulate 8 channels delivering stimulation at 250 pulses per second on each channel in a round-robin fashion (**Figure 2A right top**). Second, all neurons in any frequency band deliver the same amplitude: the average across the entire frequency band. Functionally, this is important since this degradation occurs at the afferent neural representation level, not in the sound waveform itself, posing a fundamental challenge to how previously learned cortical representations might continue to function. Yet, remarkably, many CI users are able to comprehend speech almost immediately after recovery from implantation surgery.

**Figure 2.**
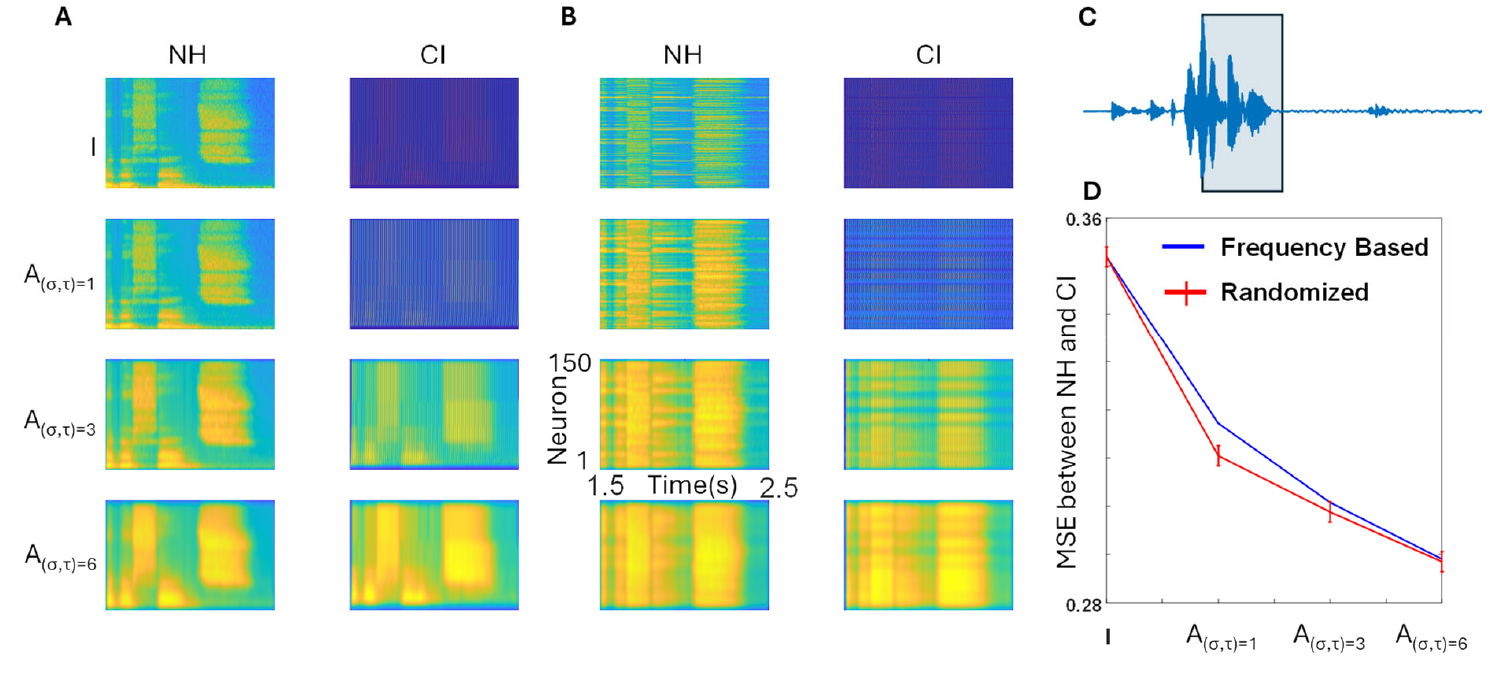
Spatiotemporal abstraction explains how cochlear implant users can recognize speech despite degraded input. (**A**) Left column: Spectrograms of a fragment of spoken sentence (“The red fox jumped over the fence”) progressively blurred using 2D Gaussian kernels with increasing spatial and temporal widths (σ = τ, from 1 to 3 to 6). These simulate hierarchical spatiotemporal abstraction layers in a normal-hearing (NH) auditory system. Each spectrogram spans 0–12 kHz on the y-axis across 150 input “neurons” logarithmically spaced by frequency, with time on the x-axis. Right column: Simulated cochlear implant (CI) input to the same network. The top panel shows an 8-channel frequency-quantized and phase-locked to implant stimulation version of the original sentence, mimicking CI processing. Lower rows show the resulting abstracted representations after STA integration, using the same Gaussian kernels as in the NH condition. (**B**) Input spectrograms from the original spectrogram (part A, NH, top row) are restructured by randomly permuting the rows—removing tonotopic organization while preserving overall content. The same permutation is applied to both the normal hearing (NH) and cochlear implant (CI) conditions. Abstraction layers then operate over these shuffled input arrangements. (**C**) The original sentence with the box outlining the example spectrogram fragment. (**D**) Mean squared error (MSE) between NH and CI representations at each abstraction level. MSE decreases monotonically as σ and τ increase, indicating that high-level abstract representations of the degraded CI input converge toward those of the original sentence. This supports the hypothesis that meaning can be preserved at coarse abstraction levels even when fine-scale input structure is lost. The same is shown for n=10 randomized spectrograms mean and std.

Consider a patient who lost hearing after acquiring normal language. Suppose they previously heard the sentence “The red fox jumped over the fence.” The auditory cortex would have received a structured spectrogram-like neural representation of this input, where neurons (plotted along the y-axis) encode frequency and time (x-axis) (**Fig. 2A, top left**). Through successive layers of STA-like processing, this input would be represented at progressively higher abstraction levels via spatiotemporal integration (**Fig. 2A, left column, lower rows**).

After cochlear implantation, the same sentence generates a sparse, quantized, and degraded input pattern (**Fig. 2A, top right**). Yet, when passed through the original STA integration layers built from normal hearing experience, the resulting abstract representations (**Fig. 2A, right column, lower rows**) closely resemble the abstracted versions of the original sentence. We quantify this similarity using mean-squared error (MSE) across abstraction layers (**Fig. 2D blue**), finding that the MSE decreases monotonically with increasing integration scale. This suggests that while the low-level signal is severely distorted, meaning can still be accessed at higher abstraction levels.

To probe the robustness of this effect, we disrupted the anatomical tonotopy by randomly shuffling the input neuron order (**Fig. 2B, top left and right**). Despite this rearrangement, the abstraction layers still preserved similarity between the NH and CI conditions (**Fig. 2D, red and blue lines**), indicating that STA does not depend on a specific spatial layout, as long as the same connections are preserved before and after distortion.

## Discussion

The cochlear implant example illustrates how STA can explain a long-standing paradox: how speech recognition remains intact even when neural input is scrambled by artificial stimulation. Despite severe distortion at the input level, abstraction layers built during normal hearing can still recover meaning by integrating over time and space. The robustness of this mechanism is demonstrated by our shuffled-tonotopy simulations, where recognition performance remains intact so long as the integration structure is preserved.

The general concepts of spatiotemporal integration and of abstraction not new. They are hallmarks of cortical processing across sensory and cognitive domains. In vision, receptive field sizes increase systematically from the retina to primary visual cortex and onward to inferotemporal areas, where neurons respond to complex objects with high tolerance to position, scale, and viewpoint changes ^3^. In audition, frequency selectivity in primary auditory cortex gives way to phonemic and linguistic encoding in higher-order areas such as the superior temporal gyrus ^14^. Similar hierarchical integration is observed in the somatosensory system ^15^, prefrontal cortex ^16^, and the entorhinal–hippocampal circuit, where place and grid cells exhibit spatial scales that vary systematically across anatomical gradients ^17^. It is important then to differentiate STA from the concepts of the conventional hierarchical abstraction, such as edge detection leading to shape identification to object encoding and categorization. Instead, STA provides robustness to the input signals via layered integration over space and time.

### STA as a Reinterpretation of Cortical Modules and Canonical Circuits

The cortex is often described as a composition of modular, layered microcircuits connected via large-scale hubs. While this architecture has long been anatomically characterized, its computational logic remains a subject of debate ^5^. We speculate that STA occurs in localized modules and provide robust, multi-scale representations via layered integration over space and time. The hierarchical abstractions would take place independent of this functionality as part of communication between modules (**Fig. 1E**).

Each local module, such as a column in V1, a patch of A1, or a hippocampal subfield can be modeled as a set of neurons with overlapping spatial and temporal receptive fields. These neurons implement a continuum of spatiotemporal filters (varying in σ and τ), producing a layered ensemble of representations that range from detailed to abstract. Although neurons in such modules often exhibit similar tuning properties when probed with an electrode, this apparent redundancy is a computational feature, not a flaw. The STA model reframes such populations as parallel, partially redundant abstraction units: they respond similarly because they each encode stable features of the input, albeit with slightly shifted integration windows. This design ensures robustness to noise and variability, a fundamental property for stable perception and memory.

Within this framework, the well-known homologous canonical cortical circuit with the repeated structure throughout the cortex consists of layered excitatory and inhibitory interconnections. Its morphological structure is well-defined and appears to be largely similar across cortex and across species, suggesting a fundamental processing scheme. Although the computational function remains enigmatic, the conventional interpretation argues for error checking and gain control through a set of feedforward and feedback connections ^18–20^. We propose that it could also be understood as the biological substrate for implementing STA kernels. Excitatory neurons with different dendritic arborizations support variable σ (spatial extent), while inhibitory subtypes sculpt τ (temporal extent). For instance, PV+ interneurons provide fast, perisomatic inhibition that constrains the onset of temporal integration, while SST+ interneurons target dendrites and support longer time constants. Recurrent excitatory connectivity further stabilizes abstract representations through recursive smoothing. In this way, the canonical circuit is not merely a wiring pattern, but a biophysical mechanism for dynamic abstraction.

At a higher level, network hubs such as those found in prefrontal, temporal, or parietal cortices, can be viewed as integrators of already-abstracted outputs from local modules. These hubs likely operate on inputs that have passed through multiple STA layers, combining them with modalities or contexts to form generalized, cross-domain representations. Crucially, the robustness of these representations depends not on precise spike timing or spatial topography, but on the stability of the abstraction hierarchy. This view helps explain how sentence meaning, object identity, or scene context can persist even when input signals are distorted, incomplete, or novel.

This reinterpretation also clarifies longstanding puzzles. For instance, why are so many neurons in a given region tuned similarly ^21–23^? Because they instantiate overlapping STA kernels. Why does cortex have such a stereotyped canonical circuit across regions? Because STA is a domain-general computational principle implemented uniformly across the brain. Why are inhibitory neurons so diverse and precisely connected ^24^? Because they dynamically control the timescales of abstraction via their modulation of τ.

In sum, STA provides a cohesive computational lens through which both local microcircuitry and global cortical organization can be understood. It offers a testable, mechanistic bridge between anatomy and function, and invites reinterpretation of canonical neuroscience principles in terms of robust, multi-scale abstraction.

### STA Interpretation of Macroscopic Neural Signals

The Spatiotemporal Abstraction (STA) framework offers a compelling explanation for several well-documented phenomena in systems neuroscience that have long appeared paradoxical under traditional assumptions.

First, local field potentials (LFPs), which reflect the summed synaptic activity of neural populations, often appear to mirror the information captured by single-unit recordings ^25^. Under the STA model, this similarity arises because both LFPs and spiking activity are filtered through overlapping spatiotemporal kernels within the same cortical module. The neurons contributing to the LFP and those firing spikes share access to the same underlying abstraction layers, differing mainly in temporal resolution and threshold. This means the apparent redundancy between these signals is a direct consequence of shared integration architecture.

Second, STA sheds light on a striking empirical observation: high-dimensional neural recordings from multielectrode arrays or calcium imaging often collapse into a small number of principal components, even during complex tasks ^26,27^. From the STA perspective, this low-dimensional structure is not surprising—neurons within a module compute overlapping abstractions of shared inputs across space and time. As a result, even large neural populations yield coordinated, low-dimensional dynamics that reflect abstracted input representations rather than raw sensory detail. These dynamics, often termed “latent variables” in dynamical systems models, may in fact correspond to the intrinsic abstraction layers of STA.

Together, these observations suggest that STA is not only a theory of local computation or perceptual robustness, but also a unifying principle that explains why neural signals across scales and recording modalities often converge onto a common low-dimensional, abstraction-preserving structure.

### Broader Implications and Hypotheses

The spatiotemporal abstraction (STA) model provides a unified computational framework for understanding how the brain preserves meaning despite degraded, variable, or incomplete input. By integrating neural activity across space and time, the brain effectively smooths fine-grained variability while retaining the structural regularities that define a concept. This layered integration enables stable, high-level representations to emerge even when low-level signals are noisy or corrupted.

To test the fundamental prediction of STA, one could record from many neurons simultaneously in a localized area of the cortex using calcium imaging. Each neuron can then be tagged as responding or not to variable changes in stimulus. Slight variations would keep the high-level integrators firing, while low-level integrators would change. This can be done for varying degrees of input to investigate the abstraction capability of any given neuron.

Beyond sensory perception, the same mechanism may underly memory encoding. By storing abstracted, multi-scale representations, cortical circuits can preserve core meaning while discarding fragile, instance-specific details. Within this context, because the highest-level abstractions (most blurred ones) in STA may behave similarly for many possible inputs, the ability to shift between attractor networks that incorporate highest levels of STA may be easier than shifting between lower-level attractors. This principle could contribute to the explanation of creativity, phantom memories, and generalizations as random fluctuations that shift network attractors between the highest-abstracted levels of STA.

Failures in this integration process due to atypical connectivity, temporal disorganization, or imbalanced emphasis on detail, could contribute to perceptual instability or cognitive deficits, positioning STA as a candidate framework for understanding certain neurological or psychiatric conditions.

The theory also informs the design of neuroprosthetic devices. Rather than reproducing high-fidelity sensory detail, successful prosthetics may instead aim to stimulate neural circuits in ways that align with the brain’s intrinsic abstraction hierarchies, optimizing interpretation rather than precision. Similarly, STA principles may inspire machine learning systems that emulate the brain’s ability to generalize from incomplete or noisy data. By embedding multi-scale, abstraction-based processing into artificial architectures, such systems could achieve greater robustness, algorithmic optimization, and interpretability in real-world environments.

In sum, STA links spatiotemporal integration to conceptual resilience, offering a biologically grounded mechanism by which meaning can survive noise. It unifies diverse findings across anatomy, physiology, and computation, and points toward a principled approach for both understanding and engineering intelligent systems. Future work should test this framework across modalities and species, develop quantitative predictions for neural encoding under degradation, and explore how manipulating abstraction hierarchies may enhance neuroprosthetic function or restore perceptual stability in clinical populations.

## References

1. Wilson, B. S. & Dorman, M. F. Cochlear implants: a remarkable past and a brilliant future. Hear Res 242, 3–21 (2008).

2. Shannon, R. V., Zeng, F. G., Kamath, V., Wygonski, J. & Ekelid, M. Speech recognition with primarily temporal cues. Science 270, 303–304 (1995).

3. DiCarlo, J. J., Zoccolan, D. & Rust, N. C. How does the brain solve visual object recognition? Neuron 73, 415–434 (2012).

4. Yamins, D. L. K. & DiCarlo, J. J. Using goal-driven deep learning models to understand sensory cortex. Nat Neurosci 19, 356–365 (2016).

5. Park, H.-J. & Friston, K. Structural and Functional Brain Networks: From Connections to Cognition. Science 342, 1238411 (2013).

6. Gollisch, T. & Meister, M. Eye Smarter than Scientists Believed: Neural Computations in Circuits of the Retina. Neuron 65, 150–164 (2010).

7. Wässle, H. & Boycott, B. B. Functional architecture of the mammalian retina. Physiol Rev 71, 447–480 (1991).

8. Field, G. D. & Chichilnisky, E. J. Information processing in the primate retina: circuitry and coding. Annu Rev Neurosci 30, 1–30 (2007).

9. Gauthier, J. L. et al. Receptive fields in primate retina are coordinated to sample visual space more uniformly. PLoS Biol 7, e1000063 (2009).

10. Segev, R., Puchalla, J. & Berry, M. J. Functional organization of ganglion cells in the salamander retina. J Neurophysiol 95, 2277–2292 (2006).

11. Kral, A. & Sharma, A. Developmental neuroplasticity after cochlear implantation. Trends Neurosci 35, 111–122 (2012).

12. Friesen, L. M., Shannon, R. V., Baskent, D. & Wang, X. Speech recognition in noise as a function of the number of spectral channels: comparison of acoustic hearing and cochlear implants. J Acoust Soc Am 110, 1150–1163 (2001).

13. Faulkner, A., Rosen, S. & Wilkinson, L. Effects of the number of channels and speech-to-noise ratio on rate of connected discourse tracking through a simulated cochlear implant speech processor. Ear Hear 22, 431–438 (2001).

14. Chang, E. F. et al. Categorical speech representation in human superior temporal gyrus. Nat Neurosci 13, 1428–1432 (2010).

15. Mountcastle, V. B. The Sensory Hand: Neural Mechanisms of Somatic Sensation. (Harvard University Press, 2005). doi:10.2307/j.ctv23dxd9k.

16. Badre, D. & D’Esposito, M. Is the rostro-caudal axis of the frontal lobe hierarchical? Nat Rev Neurosci 10, 659–669 (2009).

17. Stensola, H. et al. The entorhinal grid map is discretized. Nature 492, 72–78 (2012).

18. Harris, K. D. & Shepherd, G. M. G. The neocortical circuit: themes and variations. Nat Neurosci 18, 170–181 (2015).

19. Shipp, S. Structure and function of the cerebral cortex. Current Biology 17, R443–R449 (2007).

20. Denève, S. & Machens, C. K. Efficient codes and balanced networks. Nat Neurosci 19, 375–382 (2016).

21. Ringach, D. L. Spontaneous and driven cortical activity: implications for computation. Curr Opin Neurobiol 19, 439–444 (2009).

22. Kohn, A., Coen-Cagli, R., Kanitscheider, I. & Pouget, A. Correlations and Neuronal Population Information. Annu Rev Neurosci 39, 237–256 (2016).

23. DeWeese, M. R., Hromádka, T. & Zador, A. M. Reliability and representational bandwidth in the auditory cortex. Neuron 48, 479–488 (2005).

24. Kepecs, A. & Fishell, G. Interneuron cell types are fit to function. Nature 505, 318–326 (2014).

25. Einevoll, G. T., Kayser, C., Logothetis, N. K. & Panzeri, S. Modelling and analysis of local field potentials for studying the function of cortical circuits. Nat Rev Neurosci 14, 770– 785 (2013).

26. Cunningham, J. P. & Yu, B. M. Dimensionality reduction for large-scale neural recordings. Nat Neurosci 17, 1500–1509 (2014).

27. Churchland, M. M. et al. Neural population dynamics during reaching. Nature 487, 51– 56 (2012).

